# Cardiomyocytes Deficiency of SENP1 Promotes Myocardial Fibrosis via Paracrine Action

**DOI:** 10.1101/2023.06.21.546019

**Authors:** Zhihao Liu, Xiyun Bian, Lan Li, Li Liu, Chao Feng, Ying Wang, Jingyu Ni, Sheng Li, Dading Lu, Yanxia Li, Chuanrui Ma, Tian Yu, Xiaolin Xiao, Na Xue, Yuxiang Wang, Chunyan Zhang, Xiaofang Ma, Xiumei Gao, Xiaohui Fan, Xiaozhi Liu, Guanwei Fan

## Abstract

**Background:** SENP1 (Sentrin-specific protease 1) is involved in heart disease by regulating intracellular SUMOylation. However, the role of cell type-specific SENP1 in the setting of myocardial infarction (MI) remains to be defined. We determined whether cardiomyocyte-specific SENP1 regulates maladaptive remodeling in the ischemic heart.

**Methods:** We generated cardiomyocyte-specific SENP1 knockout and overexpression mice to assess cardiac function and ventricular remodeling responses under physiological and pathological conditions. Adult mouse cardiomyocytes and fibroblasts were separated to observe the paracrine effects of cardiomyocytes. DIA proteomic assay and bioinformatic analysis were performed to detect differential proteins in primary cardiomyocytes. Single-cell sequencing data were analyzed to identify key cardiomyocyte cluster regulated by SENP1 after MI. Mass spectrometry was performed to search for key effector proteins regulated by SENP1.

**Results:** SENP1 expression reduced in post-infarction mice and patients with heart failure. Cardiomyocyte-specific SENP1 deficiency (genetic or adeno-associated virus (AAV)-mediated) triggered a myocardial fibrotic response and progressive cardiac dysfunction. MI exacerbated the poor ventricular remodeling response mediated by SENP1 deletion, SENP1 overexpression improved cardiac function and reduced fibrosis. Further, we found that increased cardiac fibrosis in the cardiomyocyte-specific SENP1 deletion mice, associated with increased Fn expression and secretion in cardiomyocytes, promoted fibroblast activation in response to myocardial injury. Mechanistically, SENP promoted deSUMOylation of the HSP90ab1 (heat shock protein 90 alpha, class B, member 1) Lys72 and ubiquitin-dependent degradation, which reduced HSP90ab1-mediated phosphorylation of STAT3 (signal transducer and activator of transcription 3) and transcriptional activation of Fn, and then reduced fibroblast activation in the indirect coculture model. On the other hand, AAV-mediated mutation of the HSP90ab1 Lys72 ameliorated adverse ventricular remodeling and dysfunction post-MI.

**Conclusions:** SENP1 is essential for repressing STAT3-mediated Fn transcription by regulating deSUMOylation of HSP90ab1. Blockade of HSP90ab1 SUMOylation at the Lys72 site is a novel therapy effective in reversing adverse ventricular remodeling post-MI.

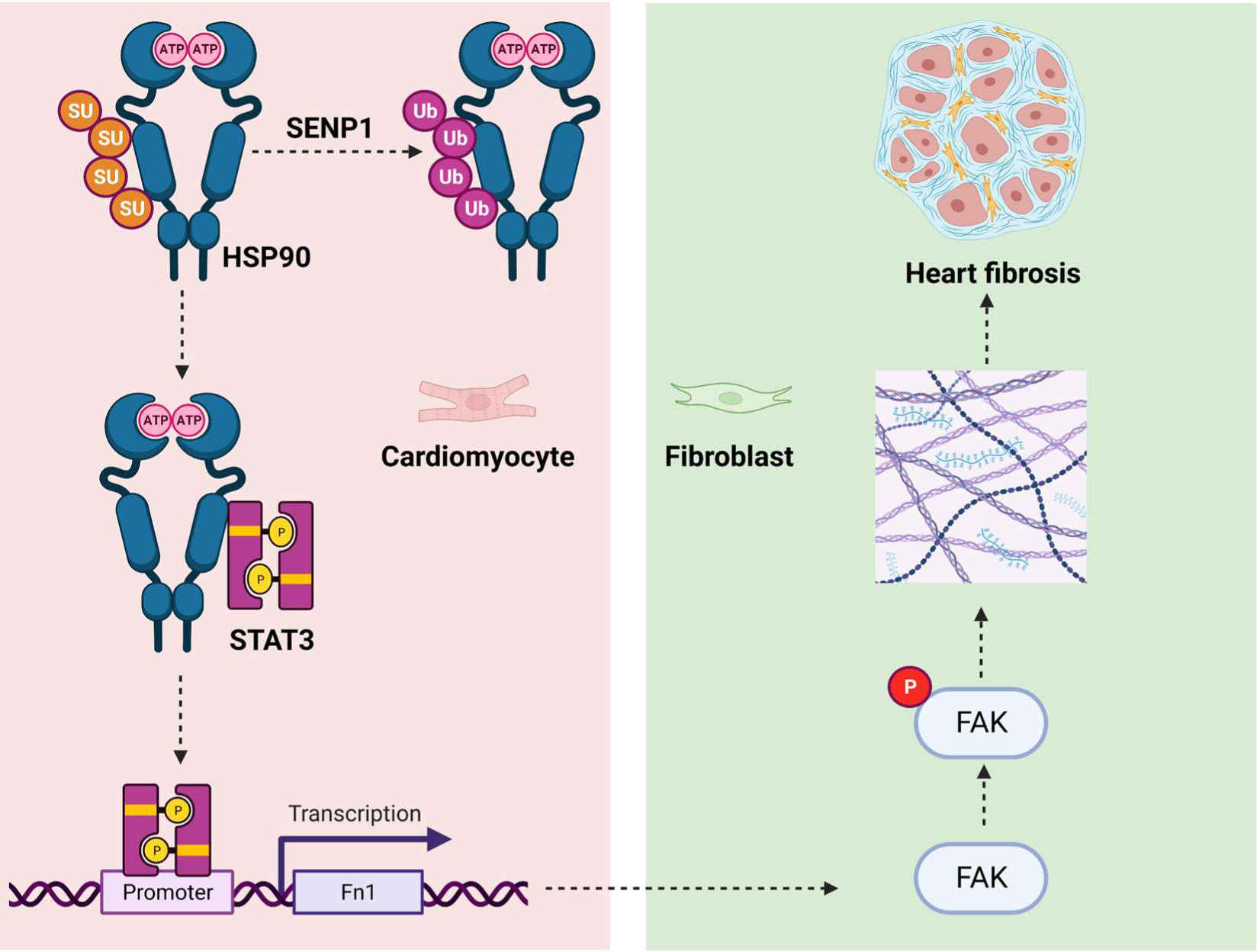

## Introduction

Myocardial infarction (MI), caused by complete or partial occlusion of coronary arteries, is one of the leading causes of death worldwide.^1^ Inflammatory cells are recruited to remove dead cardiomyocytes, and fibroblasts proliferate and secrete collagen to replace the lost cardiomyocytes. Initially, the repair response is adaptive, but excessive collagen production and scar expansion trigger a maladaptive ventricular remodeling response that promotes the progression of heart failure.^2, 3^ Therefore, understanding the molecular mechanisms responsible for the progression of maladaptive remodeling in MI is essential for the development of novel targets to reverse pathological remodeling and block the progression of heart failure.

SUMOylation is a reversible post-translational modification by covalent binding to specific lysine sites on target proteins.^4^ SUMOylation is critical for the regulation of the cell cycle, cell metabolism, gene transcription, DNA damage and repair.^5^ Normally, SUMO proteins bind to substrates through a cascade of E1-activating enzyme, E2 conjugase, and E3 ligases, while SENPs are precisely responsible for the removal of SUMO proteins from target proteins.^6^ The SENPs family consists of 6 members, SENP1, SENP2, SENP3, SENP5, SENP6, and SENP7. Of all the identified SENPs family, SENP1 was originally reported and has been extensively studied in diseases such as cancer,^7^ neurodegenerative diseases,^8^ and cerebral ischemia-reperfusion.^9^ However, the role of SENP1 in cardiovascular disease, particularly in cardiac remodeling after MI, remains unclear.

A recent study demonstrated that gene delivery system-mediated knockdown of SENP1 exacerbates pressure overload-induced myocardial hypertrophy and interstitial fibrosis, accelerating the progression of heart failure.^10^ Additionally, heterozygous SENP1 knockout aggravated myocardial ischemia/reperfusion (I/R) injury by inhibiting the expression of hypoxia-inducible factor 1α (HIF1α).^11^ These results suggest that SENP1 is an important candidate gene for the treatment of cardiovascular disease. However, previous studies have mainly focused on the whole heart to investigate the effects of SENP1, and the role of SENP1 in different types of heart cells has not been reported. Recently, we demonstrated that the regulation of pathological remodeling by SUMO after MI is cell-specific.^12^ SENP1 expression in cardiomyocytes and the direct role of SENP1 in pathological remodeling remain unknown. Therefore, it is necessary to evaluate the roles of SENP1 based on specific cell types in normal and pathological cardiac function.

In this study, we not only found that SENP1 was significantly decreased in the post-infarct hearts but also generated a cardiomyocyte-specific SENP1 inducible KO mouse, which exhibited markedly increased ischemia-induced fibrotic response. Unexpectedly, we found that increased myocardial fibrosis in cardiomyocytes-specific SENP1 deletion mice was associated with enhanced Fn secretion and upregulation of HSP90ab1 in vitro and in vivo. Finally, we demonstrate that inhibiting HSP90ab1 SUMOylation using a gene intervention strategy in cardiomyocytes improves cardiac function after MI as a therapeutic intervention.

## Methods

### Data Availability

All supporting data are available within the article and in the Data Supplement. Detailed descriptions of the experimental methods are provided in the Data Supplement.

## Results

### SENP1 Expression Is Downregulated in Cardiomyocytes During Cardiac Remodeling Progression

To determine whether SENP1 is involved in the pathological remodeling response, we examined the expression of SENP1 at different time points after MI in mice. In an injury-time course, we found that SENP1 expression decreases already at 5 days post-MI, and it is still downregulated at 1 week although to a significantly lower extent (Figure 1A and 1B). In addition, we collected biopsy samples from 4 patients subjected to a trans jugular interventricular septal myocardial biopsy and classified the patient’s heart function according to NYHA (Online Table I). Immunoblot analysis showed that SENP1 protein levels were also dramatically reduced in the hearts of patients diagnosed with severe heart failure (Figure S1A and S1B). To explore the distribution of SENP1 in cardiac tissue, we performed a single-cell transcriptome analysis in our previously characterized mouse model of MI after coronary artery ligation.^12^ Unbiased single-cell transcriptomic analysis revealed a significant reduction in SENP1 levels in cardiomyocytes at 7 days after MI (Figure 1C). Consistent with the single-cell transcription data, higher SENP1 protein levels were observed in cardiomyocytes compared with fibroblasts and endothelial cells (Figure S1C), suggesting a role for cardiomyocyte SENP1 during cardiac remodeling.

**Figure 1.**
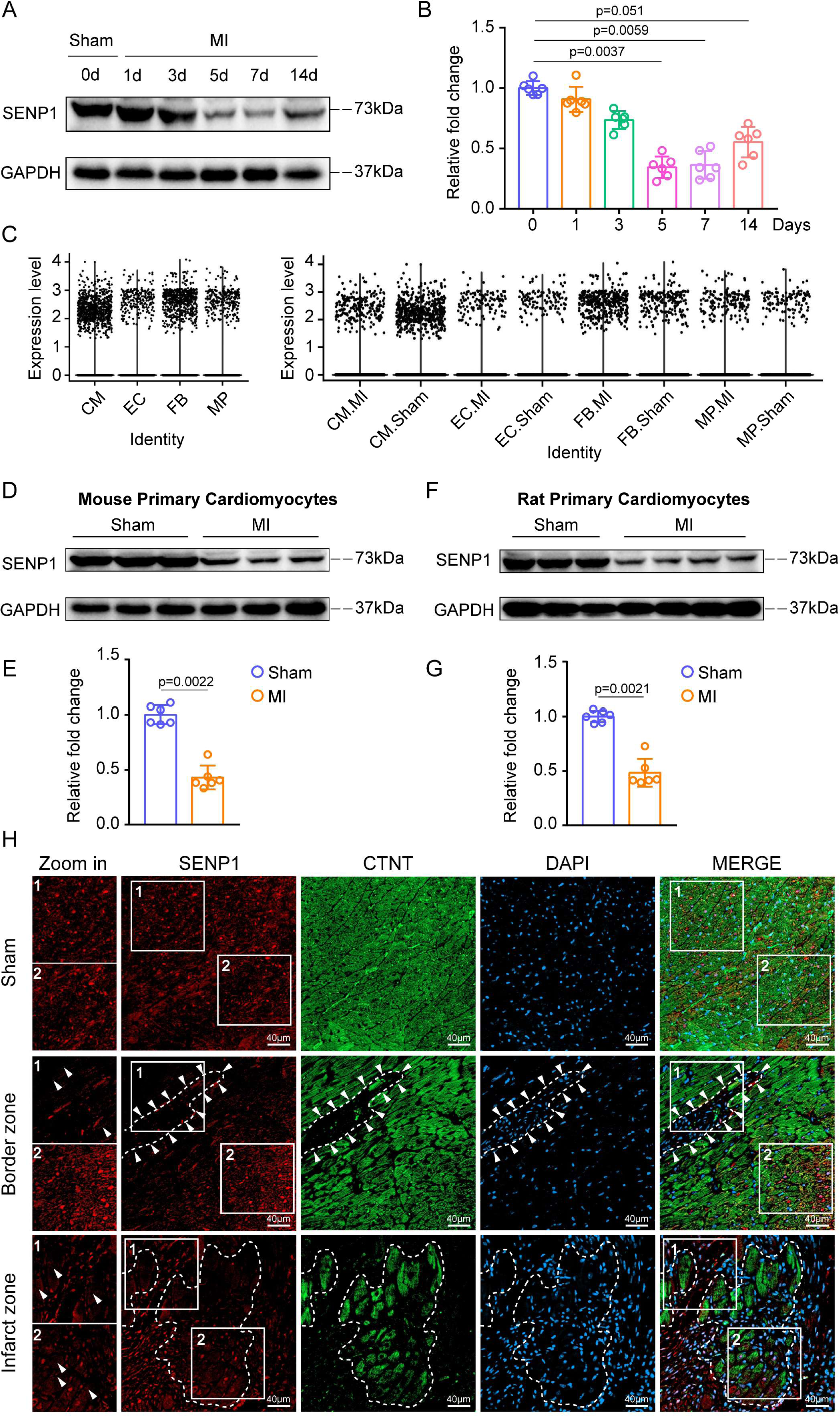
Myocardial injury triggers downregulation of SENP1 in cardiomyocytes. **A and B,** Time course of SENP1 expression in response to myocardial injury with representative immunoblot images in **A**. Total proteins were extracted and subjected to analysis of SENP1 protein levels. Quantitative analyses were plotted in **B**; n=6 in each group. **C,** Violine plot showing SENP1 scaled expression across the whole cell types in the mouse heart snRNA-seq data. **D and E,** Protein levels of SENP1 in mouse primary cardiomyocytes after MI with representative immunoblot images in **D** and quantitative analyses were plotted in **E**; n=6 in each group. **F and G,** Protein levels of SENP1 in rat primary cardiomyocytes after MI with representative immunoblot images in **F** and quantitative analyses were plotted in **G;** n=6 in each group. **H,** Immunofluorescent staining of SENP1 (red) and CTNT (green) was performed in sham and MI-injured mouse heart. One-way ANOVA followed by the Dunn post hoc multiple comparisons test was conducted in **B**. Mann-Whitney U test was utilized in **E** and **G.**

The expression level of SENP1 was detected by primary mice ventricular myocytes after MI, a lower SENP1 protein level was observed after myocardial injury than healthy (Figure 1D and 1E). In addition to mice, the downregulation of SENP1 expression post-MI was also observed in rats (Figure 1F and 1G). As shown in Figure S1D through S1F, SENP1 was significantly decreased in NRVMs in response to ischemic stimulation. To confirm the involvement of SENP1 in cardiomyocyte induction by myocardial injury, we performed immunofluorescence analysis of SENP1 protein levels in cardiac cross-sections from mice after myocardial injury. Similar to the decrease in SENP1 protein levels and distribution shown in our myocardial injury model, we found reduced SENP1 in the cardiomyocytes surrounding the infarct area (not the distal region) injured by 5 days of MI (Figure 1H). Importantly, we also found reduced SENP1 expression in cardiomyocytes from patients with severe heart failure (Figure S1G). Collectively, these results showed that SENP1 in myocytes is a critical source in response to myocardial injury and that SENP1 may play a role in cardiac remodeling.

### Loss of SENP1 in Cardiomyocytes Induces Cardiac Dysfunction and Myocardial Fibrosis

To assess the role of SENP1, we selectively deleted the SENP1 gene in cardiomyocytes. SENP1^flox/flox^ mice were bred with CRE recombinase expressed under myosin heavy chain 6 (Myh6-Cre) transgenic mice. Cardiomyocytes and non-cardiomyocytes were separated by Langendorff perfused isolated mouse hearts and homogenized to detect SENP1 protein levels (Figure S2A). Eight-week-old mice were measured by ultrasound from 1 to 4 weeks after injection of tamoxifen (TAM) 5 days. We found that cardiac performance was significantly reduced in SENP1^flox/flox^Myh6^Cre^ mice compared to controls after tamoxifen induction, including reduced ejection fraction and fractional shortening. The observed cardiac dysfunction persisted for 4 weeks after cardiomyocyte-specific SENP1 knockout (Figure 2A and 2B). Histological analysis by Masson’s trichrome staining exhibited obvious evidence of interstitial fibrosis within the ventricular myocardium after SENP1 deletion (Figure 2C and 2D). Significant deposition of collagen was observed in the SENP1^flox/flox^Myh6^Cre^ mice as shown by immunohistochemical staining and immunofluorescence (Figure 2E through 2H). In addition, we observed significantly increased expression of fibrotic protein, including collagen 1 and α-SMA, which were significantly elevated in the SENP1flox hearts (Figure 2I and 2J). This deleterious response was not associated with differences in cardiomyocyte hypertrophy or apoptosis following cardiomyocyte-specific SENP1 loss. (Figure S2B through S2E).

**Figure 2.**
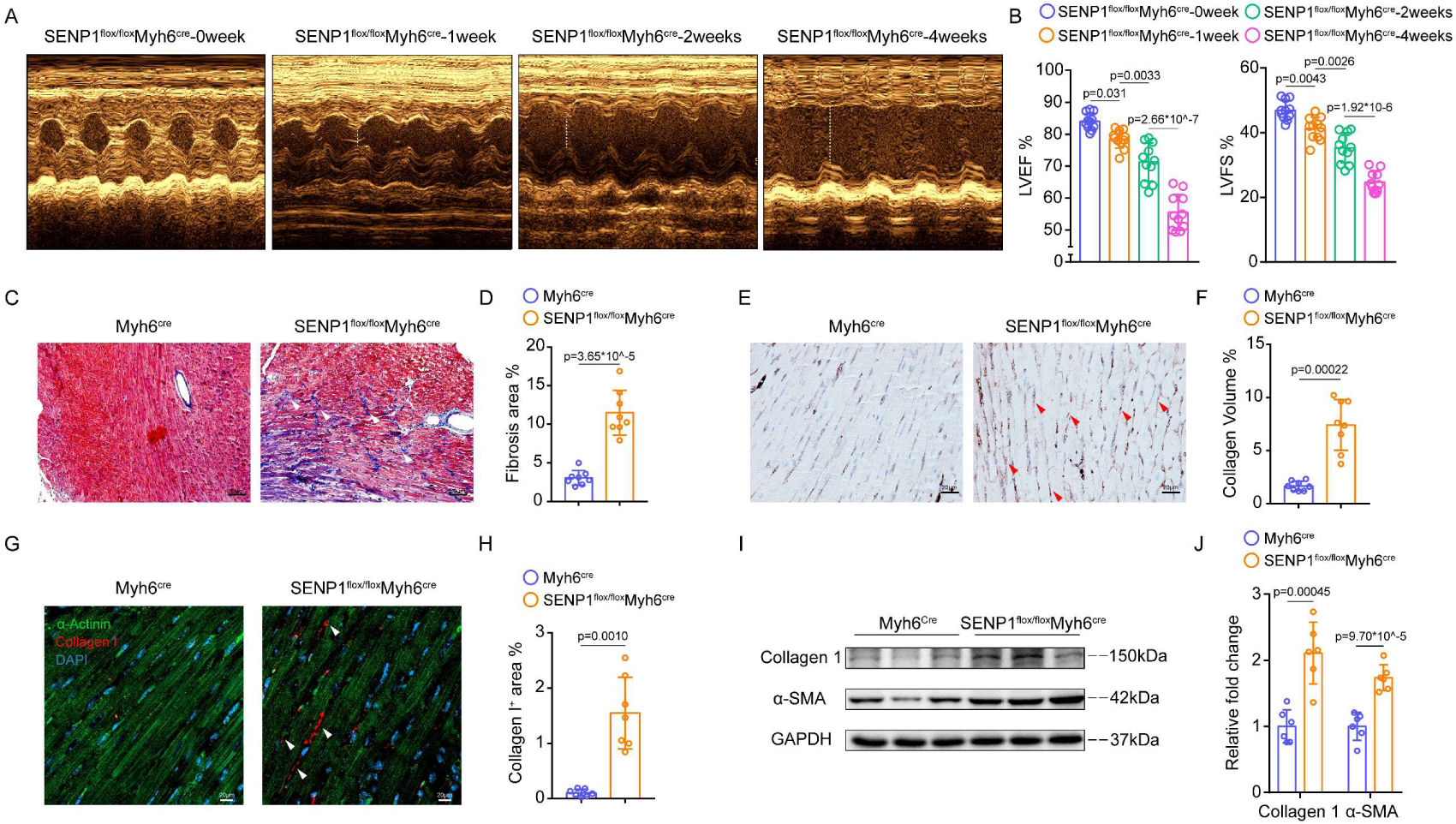
Loss of SENP1 in cardiomyocytes incurs cardiac fibrosis accompanied by deteriorated cardiac function. **A,** Representative echocardiographic images of SENP1^flox/flox^Myh6^cre^ mice. **B,** LVEF and LVFS measured by 2-dimensional echocardiography on mice; n=11 in each group. **C and D,** Representative images of left ventricle stained with Masson’s trichrome staining in **C**; scale bar, 50 μm; Quantitative analyses were plotted in **D**; n=8 in each group. **E and F,** Representative images of left ventricle stained with Collagen I staining in **E**; scale bar, 20 μm; Quantitative analyses were plotted in **F**; n=8 in each group. **G and H,** Immunofluorescence staining for collagen I^+^ (red) and α-actin^+^ (green) cardiac sections in **G**; scale bar, 20 μm; Quantitative analyses were plotted in **H**; n=7 in each group. **I and J,** Immunoblots for collagen 1 and α-SMA in SENP1^flox/flox^Myh6^cre^ and Myh6^cre^ mice heart tissues **(I).** Quantitative analyses are plotted in **J.** n=8 in each group. One-way ANOVA followed by Tukey post hoc multiple comparisons test was conducted in **B.** Unpaired t test with Welch’s correction was conducted in **D, F, H** and **J.**

To further clarify the functional significance of SENP1 in myocardial fibrosis, we performed tail vein injection of miR30-based SENP1 shRNA (CtnT promoter) (1×10^11 v.g/ml, 100μl/animal) in mice. The efficacy of the shRNA-mediated SENP1 defection was evaluated at protein level by immunoblot (Figure S3A). Echocardiography reveled a progressive reduction in ejection fraction and fractional shortening in SENP1 shRNA-treated mice (Figure S3B and S3C). Histological staining of heart sections showed the expansion of myocardial fibrosis when SENP1 was silenced (Figure S3D through S3I). Meanwhile, the expression levels of collagen 1 and α-SMA were significantly increased in shRNA-mediated SENP1-deficient hearts (Figure S3J and S3K). These results showed that SENP1 deletion in cardiomyocytes leads to myocardial fibrosis and progressive cardiac dysfunction.

### Loss of SENP1 Exacerbates Cardiac Dysfunction and Fibrosis Following Acute Myocardial Injury

To test whether SENP1 plays an important role in regulating cardiac remodeling and function after myocardial injury, we measured how the heart responds to cardiac ischemia in the absence of SENP1 in cardiomyocytes by independently assessing cardiac function at 1 to 14 days post-MI (Figure 3A). Strikingly, cardiomyocyte specific SENP1-dedicient mice showed accelerated the damage of cardiac function post-MI (Figure 3B and 3C). Since we have shown that deletion of SENP1 in cardiac myocytes establishes profibrotic functions under physiological conditions, we reasoned that SENP1 may be involved in the development of cardiac fibrosis post-MI and the subsequent development of cardiac dysfunction. A significantly increased fibrosis area and α-SMA^+^ area around cardiac myocytes were observed in SENP1^flox/flox^Myh6^Cre^ mice compared to Myh6^Cre^ mice after myocardial injury (Figure 3D through 3G). In addition, western blot analysis revealed significantly increased expression of the fibrotic proteins after SENP1 deletion, including collagen I, periostin and α-SMA (Figure 3H and 3I). Similarly, loss of SENP1 in cardiomyocytes using adenoviral strategies promoted the development of myocardial fibrosis post-MI (Figure 3J and 3K). Notably, cardiomyocyte apoptosis was exacerbated in SENP1^flox/flox^Myh6^Cre^ mice heart cross-sections after MI, compared to the control group (Figure S4), indicating that silencing of SENP1 in cardiac myocytes exacerbated adverse cardiac remodeling after ischemic injury, defined as severely impaired cardiac function, exacerbated myocardial fibrosis, and increased cardiomyocyte apoptosis.

**Figure 3.**
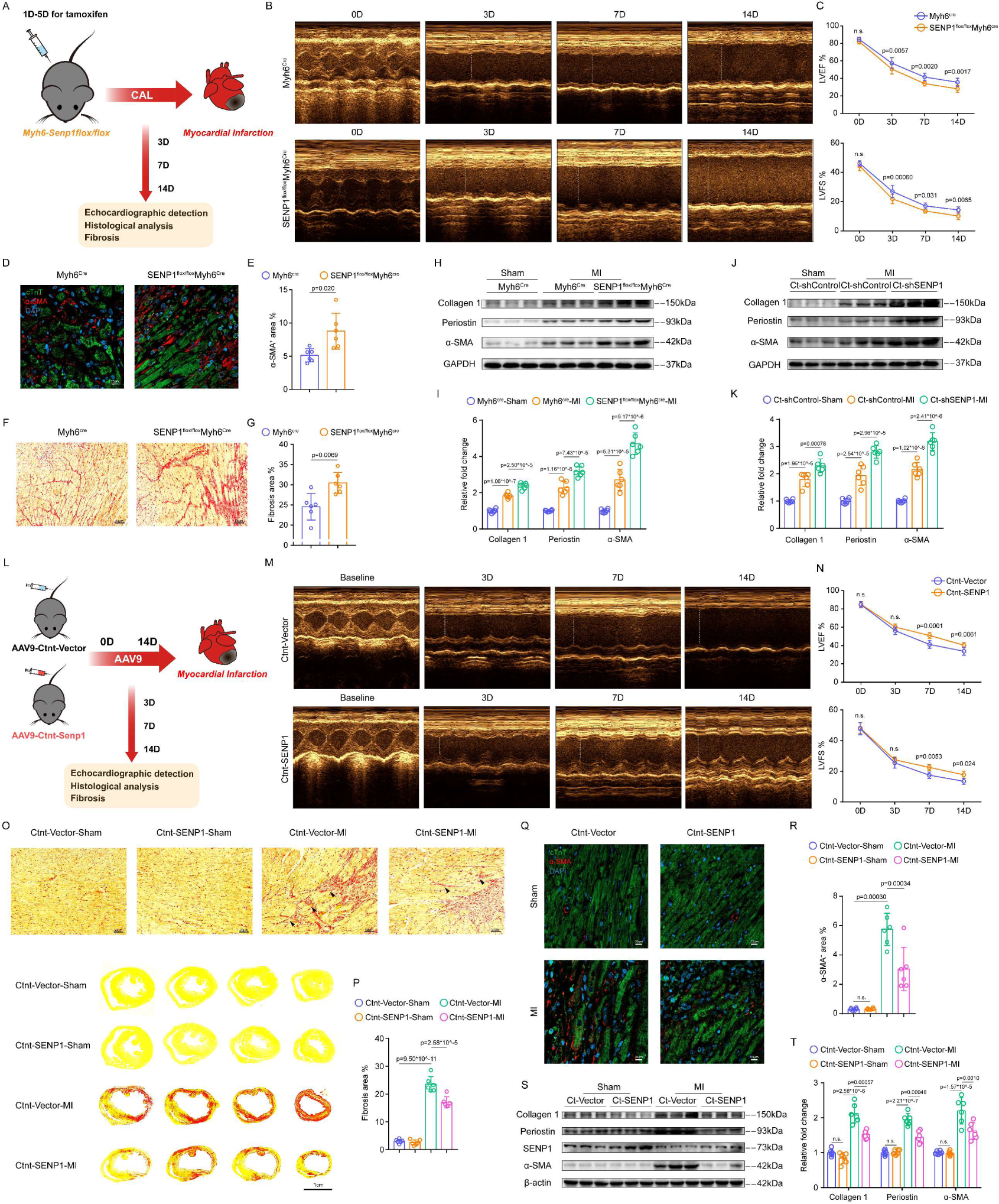
SENP1 deficiency in cardiomyocytes exacerbates fibrosis after myocardial injury and SENP1 overexpression ameliorates adverse ventricular remodeling. **A,** Schematic of MI model and echocardiography in SENP1^flox/flox^Myh6^cre^ and Myh6^cre^ mice. **B,** Representative echocardiographic images of SENP1^flox/flox^Myh6^cre^ mice and Myh6^cre^ mice after MI. **C,** LVEF and LVFS measured by 2-dimensional echocardiography on mice, at different time points; n=9 in each group. **D,** Immunofluorescence staining for α-SMA^+^ (red) and CTNT^+^ (green) cardiac sections; scale bar, 20 μm. **E,** Quantification of α-SMA^+^ areas per field; n=6 in each group. **F and G,** Representative images of left ventricle stained with Sirius red staining showing degrees of fibrosis in **F**; scale bar, 50 μm; Quantitative analyses were plotted in **G**; n=6 in each group. **H and I,** Immunoblots for collagen 1, periostin, and α-SMA in SENP1^flox/flox^Myh6^cre^ and Myh6^cre^ mice heart tissues after MI **(H).** Quantitative analyses are plotted in I. n=6 in each group. **J and K,** Immunoblots for collagen 1, periostin, and α-SMA in CTNT-shSENP1 and CTNT-shControl mice heart tissues after MI **(J).** Quantitative analyses are plotted in **K.** n=6 in each group. **L,** Schematic of MI model and echocardiography in CTNT-SENP1 and CTNT-Vector mice. **M,** Representative echocardiographic images of CTNT-SENP1 mice and CTNT-Vector mice after MI. **N,** LVEF and LVFS measured by 2-dimensional echocardiography on mice, at different time points (0, 3, 7, and 14 days after MI); n=7 in each group. **O and P,** Representative images of left ventricle stained with Sirius red staining showing degrees of fibrosis in mice; scale bar, 50 μm **(top)** and 1 cm **(bottom)**. **P,** Quantitative analyses were plotted in **P**; n=6 in each group. **Q and R,** Immunofluorescence staining for α-SMA^+^ (red) and CTNT^+^ (green) cardiac sections in **Q**; scale bar, 20 μm; Quantification of α-SMA^+^ areas per field in **R**; n=6 in each group. **S and T,** Immunoblots for collagen 1, periostin, and α-SMA in CTNT-SENP1 and CTNT-Vector mice heart tissues after MI **(S).** Quantitative analyses are plotted in **T.** n=6 in each group. Two-way ANOVA with Sidak’s multiple comparisons test was conducted in **C** and **N.** Unpaired t test with Welch’s correction was conducted in **E** and **G.** One-way ANOVA with Tukey’s multiple comparisons test was conducted in **I** and **K.** Two-way ANOVA followed by Sidak post hoc multiple comparisons test was conducted in **R, P** and **T.**

### Genetic Overexpression of SENP1 in Cardiomyocytes Reduces Adverse Ventricular Remodeling Progression

To further determine the critical role of SENP1 in the pathogenesis of pathological myocardial remodeling in vivo, recombinant adeno-associated virus (serotype 9) vectors carrying mouse SENP1 (accession number NM_144851) with a c-TNT promoter (AAV9-CTNT-SENP1) were used to investigate the effect of SENP1 overexpression in cardiomyocytes on the degree of myocardial fibrosis after MI (Figure 3L). Non-invasive echocardiography was used to determine the cardiac function and quantified by percent left ventricular ejection fraction and fractional shortening. Ventricular function was significantly improved in cardiomyocyte-specific SENP1 overexpression mice as compared to control mice 3 to 14 days post-MI (Figure 3M and 3N). Fibrotic area was significantly reduced in SENP1 overexpression mice as compared to control mice (Figure 3O and 3P). In addition, less α-SMA^+^ area was found adjacent to the interstitial space in the boundary regions after overexpression of SENP1 in cardiomyocytes (Figure 3Q and 3R). Moreover, molecular markers of myocardial fibrosis in vivo were strongly suppressed by SENP1 overexpression at both protein and mRNA levels (Figure 3S and 3T, Figure S5). Taken together, loss of SENP1 in cardiomyocytes promoted increased cardiac fibrosis and exacerbated cardiac dysfunction following acute myocardial injury in mice, and overexpression of SENP1 in cardiomyocytes inhibited excessive myocardial fibrosis and adverse ventricular remodeling.

### The Paracrine Effect of Cardiomyocytes-Fibroblasts Is Regulated by SENP1

The observation that SENP1 knockout mice have caused cardiac fibrosis and functional decompensation under physiological conditions. However, we found no apoptotic response in cardiomyocytes, which prompted us to test whether cardiomyocytes, upon deletion of SENP1 induction, would signal directly to cardiac fibroblasts. To address these questions, we isolated primary cardiomyocytes from Myh6^Cre^ and SENP1^flox/flox^Myh6^Cre^ mice and performed intervention under normal and ischemic (hypoxia and serum starvation) conditions. We then transferred mouse primary fibroblasts or embryonic fibroblast cell lines (NIH-3T3) into cardiomyocyte media (Figure 4A). Immunofluorescence analysis showed a significantly increased number of activated (α-SMA^+^) and proliferative (EDU^+^) fibroblasts in SENP1-deficient primary myocyte medium (Figure 4B through 4G). To further test this hypothesis, we cultured cardiac fibroblasts with the conditioned culture medium obtained from cardiomyocytes infected with AAV-mediated deletion of SENP1. As expected, treatment of cardiac fibroblasts with the conditioned medium for 24 hours significantly increased fibroblast proliferation and myofibroblast differentiation compared to shControl medium, and this effect was enhanced by pre-ischemia of cardiomyocytes (Figure 4H and 4I, Figure S6A and S6B). Moreover, molecular markers of myocardial fibrosis in mouse primary fibroblasts were strongly promoted by SENP1-deficient primary myocyte medium at both mRNA and protein levels (Figure 4J through 4L). These results demonstrate that SENP1 deletion in cardiomyocytes accelerates fibroblast proliferation and activation through a paracrine effect.

**Figure 4.**
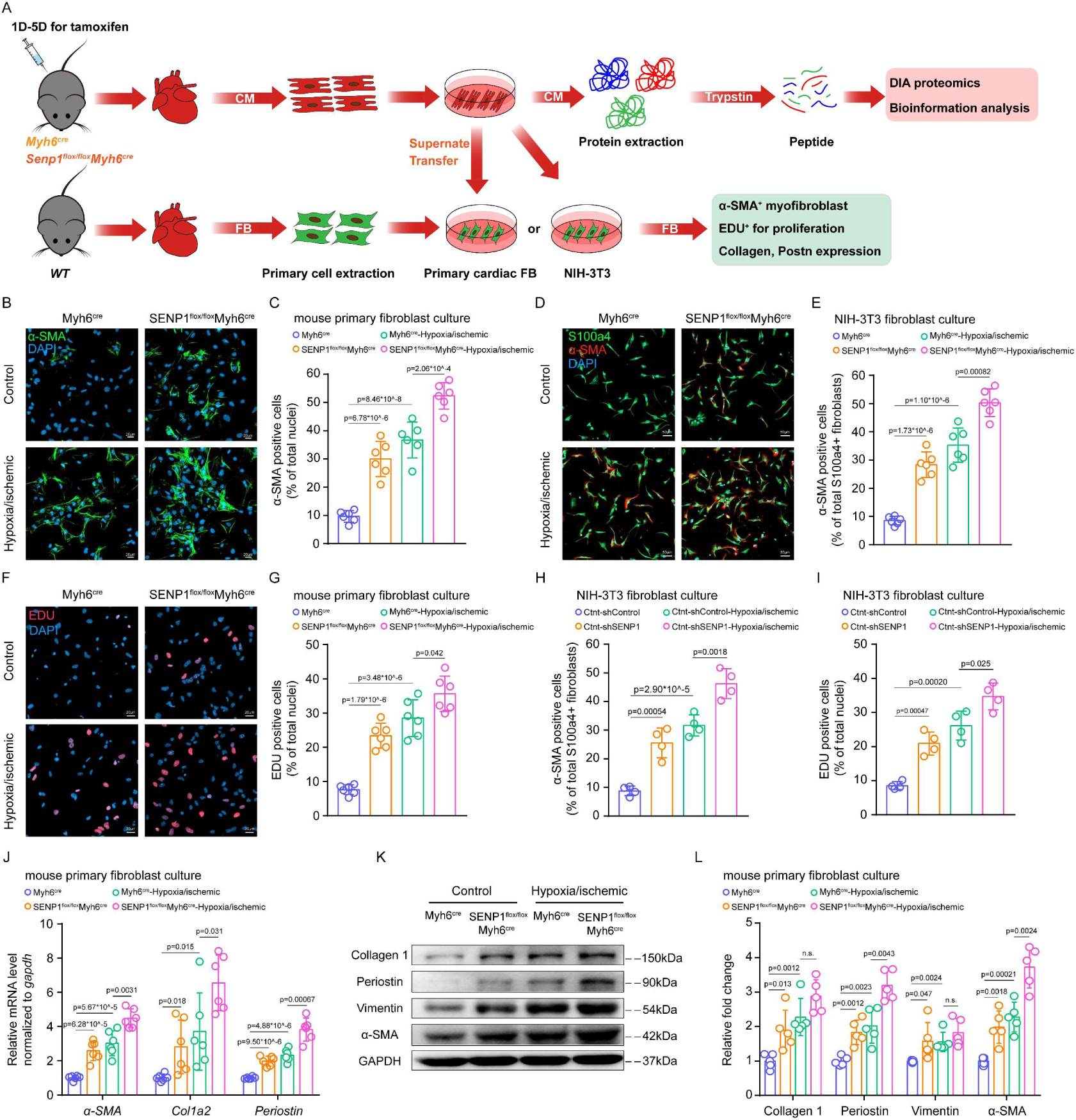
Conditioned medium collected from cardiomyocytes deficient in SENP1 increases the differentiation and proliferation of fibroblasts. **A,** Schematic representation of the in vitro assay for paracrine signaling between primary cardiac myocytes and primary fibroblasts or NIH-3T3. Conditioned medium derived from SENP1^flox/flox^Myh6^cre^ and Myh6^cre^ mice primary cardiomyocytes for fibroblast stimulation and primary cardiomyocytes for DIA proteome sequencing. **B,** Immunofluorescence staining for α-SMA^+^ (green) primary fibroblasts; scale bar, 20 μm. **C,** Quantification of α-SMA^+^ cells per field; n=6 in each group. **D,** Immunofluorescence staining for S100a4^+^ (green) and α-SMA^+^ (red) NIH-3T3 fibroblasts; scale bar, 50 μm. **E,** Quantification of S100a4-α-SMA double positive cells per field; n=6 in each group. **F,** Immunofluorescence staining for EDU^+^ (red) primary fibroblasts; scale bar, 20 μm. **G,** Quantification of EDU^+^ cells per field; n=6 in each group. **H and I,** Quantification of S100a4-α-SMA double positive cells per field **(H)** and EDU^+^ cells per field **(I) o**f NIH-3T3 fibroblasts. n=6 in each group. **J,** *α-SMA*, *Col1a2*, and *periostin* mRNA expression in mouse primary fibroblasts stimulated by conditioned medium; n=6 in each group. **K and L,** Immunoblots for collagen 1, periostin, vimentin, and α-SMA in mouse primary fibroblasts after conditioned medium stimulation **(K).** Quantitative analyses are plotted in **L.** n=5 in each group. Two-way ANOVA followed by Sidak post hoc multiple comparisons test was conducted in **C, E, G, H, I, J,** and **L.**

### Deletion of SENP1 Promotes Fn Expression and Secretion in Cardiomyocytes

To identify putative paracrine factors, primary cardiomyocytes from Myh6^Cre^ and SENP1^flox/flox^Myh6^Cre^ mice were subjected to DIA assay and bioinformatics analysis. We found that the secreted protein fibronectin (Fn) was significantly increased in SENP1-deficient cardiomyocytes (Figure 5A and 5B, Online Table II and Table III). Fn is a highly conserved protein with multiple biological roles in development, cell growth and differentiation, adhesion, migration, and wound healing, mainly through integrin-mediated signaling.^13^ Moreover, cardiac myocytes in shSENP1 group had higher Fn mRNA and protein levels than those in the shControl group (Figure 5C through 5E). We then used immunoblotting to detect Fn in the cell lysate and conditioned medium, and found that the absence of SENP1 promoted the expression and release of Fn in primary cardiac myocytes, and this effect was enhanced by pre-ischemia of the cardiac myocytes. However, Fn expression was significantly reduced in myocytes overexpressing SENP1 compared to ischemic myocytes (Figure 5F). To investigate the expression status of Fn in pre- and post-infarction mouse hearts, we then analyzed single-cell RNA sequencing data from cardiomyocytes alone. Consistent with the data in Figure 5C, *Fn* expression was increased in cardiomyocytes subsets after MI, particularly in the CM-3 cluster characterized by *Nppa*, which responds to myocardial injury (Figure S7A through S7C). Recombinant Fn was administered to primary mouse fibroblasts, and the pro-fibrosis effects of Fn were found to be concentration dependent (Figure 5G). Given that focal adhesion kinases (FAKs) are the most prominent proteins involved in the ECM-integrin signaling pathways critical for fibroblast proliferation/activation,^14^ we used a neutralizing antibody (Fn Ab) and a selective FAK inhibitor (PF-573228) to detect the activation of FAK signaling. Importantly, application of Fn Ab and PF-573228 in the medium reduced the abundance of phospho(p)-FAK, collagen I, periostin and α-SMA in fibroblasts caused by the absence of SENP1 in cardiomyocytes (Figure 5H through 5J). Together, Fn was released from damaged cardiomyocytes and mediated fibroblast proliferation and activation, and the loss of SENP1 exacerbated the production and release of Fn in cardiomyocytes.

**Figure 5.**
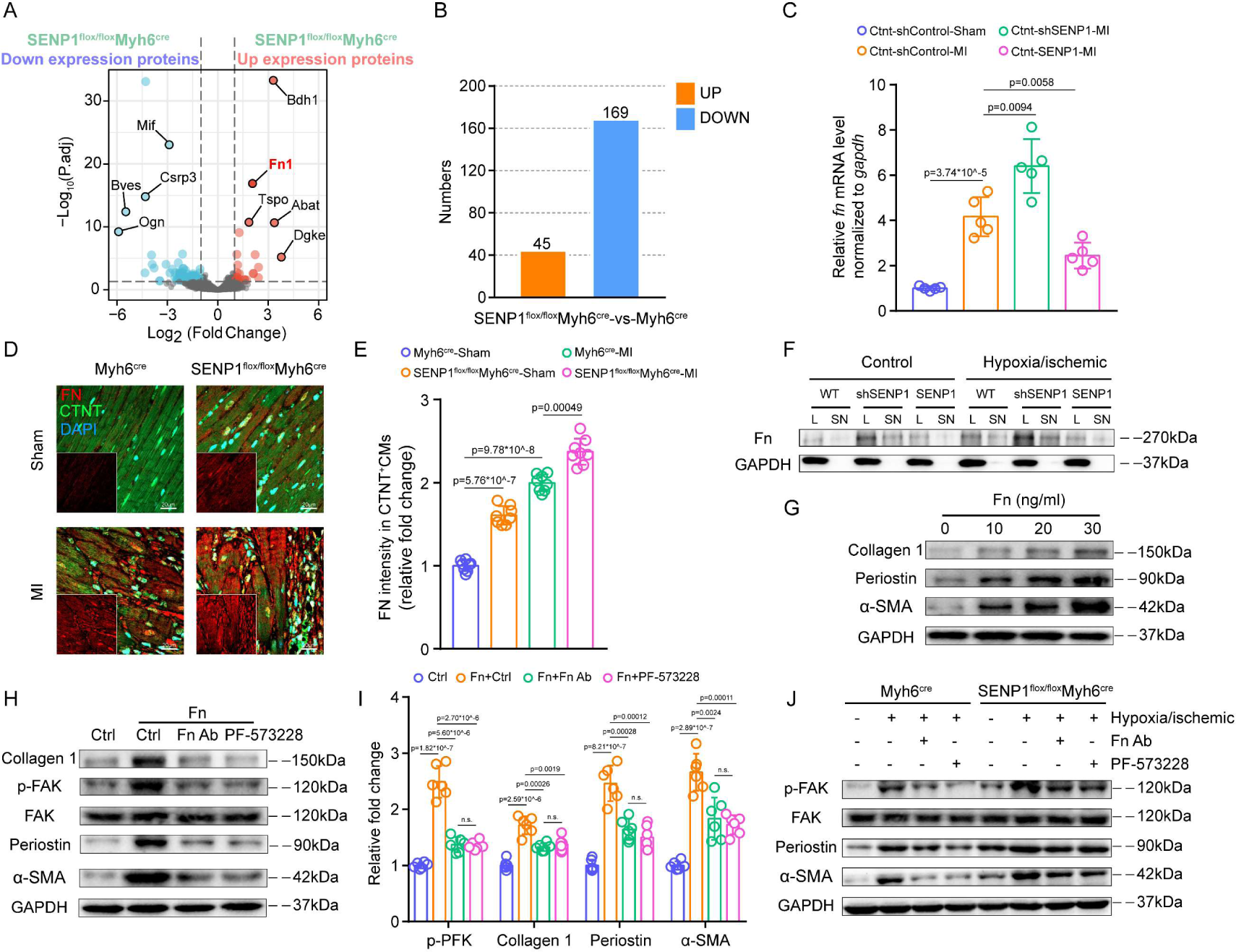
Loss of SENP1 in cardiomyocytes promotes Fn secretion and activation of FAK signaling in fibroblasts. **A,** Volcano plot of differentially expressed proteins in SENP1^flox/flox^Myh6^cre^ primary cardiomyocytes compared with those in Myh6^cre^ primary cardiomyocytes. **B,** Number of differential proteins in SENP1^flox/flox^Muh6^cre^ primary cardiomyocytes compared with Myh6^cre^ primary cardiomyocytes. **C,** *Fn* mRNA expression in mouse primary cardiomyocytes; n=5 in each group. **D,** Immunofluorescence staining for Fn (red) and CTNT (green) cardiac sections; scale bar, 20 μm. **E,** Quantification of Fn^+^ areas per field; n=8 in each group. **F,** Immunoblots for Fn in mouse primary cardiomyocytes and conditioned medium from adenovirus-transfected mice. **G,** Immunoblots for collagen 1, periostin, and α-SMA in primary fibroblasts cultured in the absence (control) or presence of Fn (at indicated concentrations). **H and I,** Immunoblots for collagen 1, periostin, and α-SMA in primary fibroblasts cultured in the presence of Fn neutralizing antibody and PF-573228 (inhibitors of FAK signaling) **(H).** Quantitative analyses are plotted in **I.** n=6 in each group. **J,** Immunoblots for FAK, p-FAK, periostin, and α-SMA, in primary fibroblasts cultured in the presence of Fn neutralizing antibody and PF-573228 after conditioned medium stimulation. One-way ANOVA with Tukey’s multiple comparisons test was conducted in **C** and **I.** Two-way ANOVA followed by Sidak post hoc multiple comparisons test was conducted in **E.**

### SENP1 Inhibits STAT3-Mediated Fn Transcription in Cardiomyocytes

We used the GTRD database (http://gtrd20-06.biouml.org/), the hTFtarget database (http://bioinfo.life.hust.edu.cn/hTFtarget#!/), and the HumanTFDB database (binding score >21) (http://bioinfo.life.hust.edu.cn/HumanTFDB#!/) to screen for transcription factors (TFs) at the *Fn* promoter and identified 14 TFs, followed by sequence alignment using JASPAR (https://jaspar.genereg.net/) to verify motifs. We found that FXOA2, IRF1, SRF, STAT3 and SP1 have motifs that bind to the *Fn* promoter, with STAT3 having the strongest site-specific binding (Figure S8A, Online Table IV). In addition, published ChIP-seq data reveled 19 STAT3 binding sites within the 1kb region of the *Fn* TSS, suggesting that the potential for direct transcriptional regulation of Fn by STAT3.^15^ Conserved binding sequences in the promoter regions of *Fn* were identified using JASPAR (https://jaspar.genereg.net/) (Figure 6A). CHIP analysis showed that STAT3 binds to the *Fn* promoter after myocardial injury, and this was enhanced by SENP1 deletion. However, overexpression of SENP1 in myocytes inhibited the occupancy of STAT3 in the *Fn* promoter region (Figure 6B and 6C). Western blot analysis was performed on infarct border and distal myocytes of mice, STAT3 Tyr-705 phosphorylation and nuclear STAT3 content were significantly increased after MI, especially in the border region (Figure 6D and 6E). Western blot and immunofluorescence assays showed that myocardial injury increased STAT3 phosphorylation and nuclear translocation. As expected, loss of SENP1 strongly increased STAT3 phosphorylation and nuclear translocation in cardiac myocytes, which was blocked by overexpression of SENP1 (Figure 6F through 6I). We further investigated whether STAT3 inhibition attenuated the intense pro-fibrosis mediated by SENP1 deletion in cardiomyocytes. Immunofluorescence analysis revealed a significantly reduced number of activated fibroblasts in SENP1flox primary myocyte medium after application of Stattic compared to SENP1flox primary myocyte medium (Figure 6J and 6K). These results demonstrate that SENP1 inhibits STAT3 activation and reduces myocardial injury–induced Fn transcription in cardiomyocytes.

**Figure 6.**
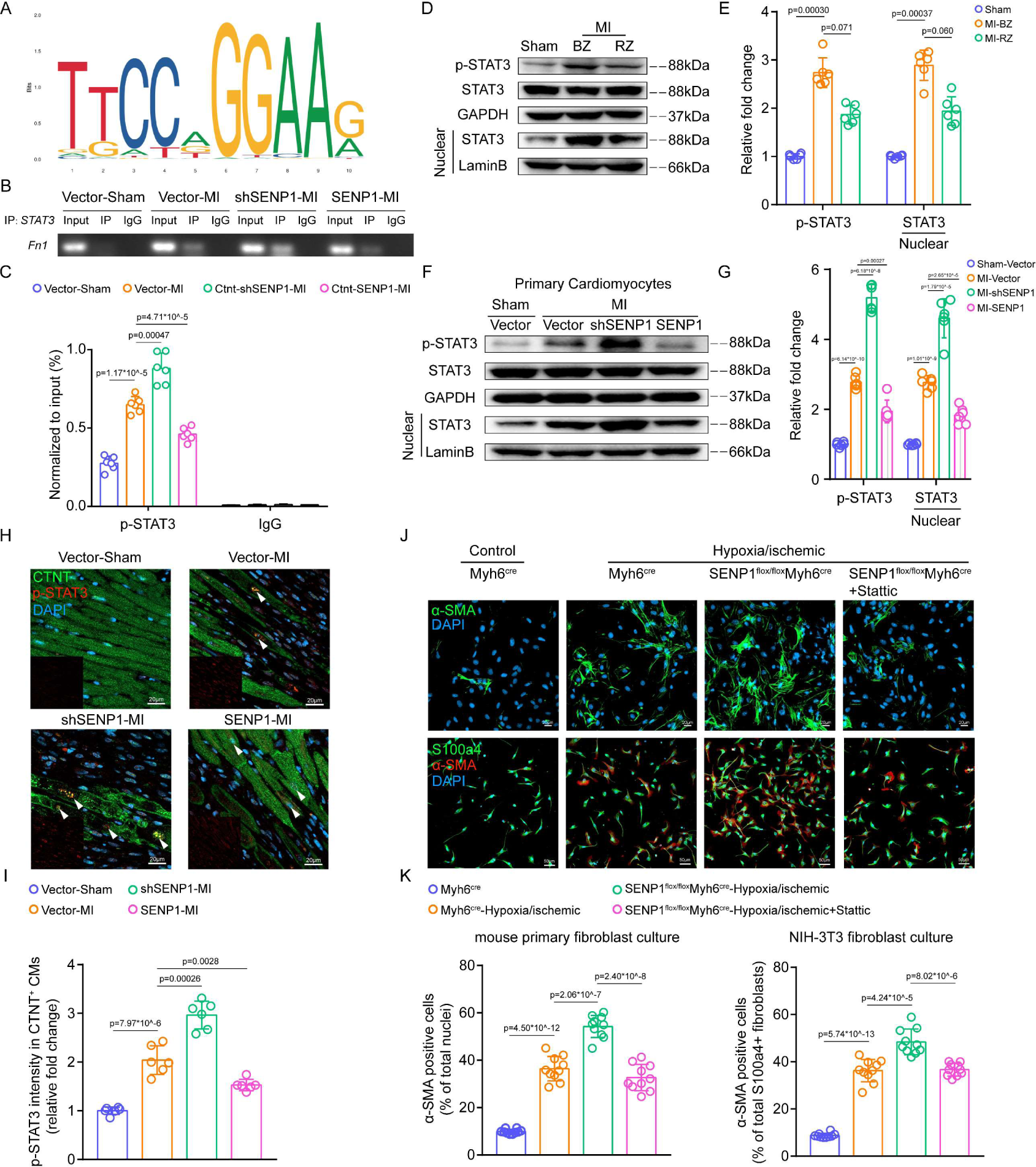
SENP1 represses STAT3-mediated transcription of Fn in cardiomyocytes. **A,** Schematic representation of conserved STAT3 binding sequences identified in the promoter regions of *Fn* using JASPAR. **B and C,** The DNA fragment derived from mouse primary cardiomyocytes corresponding to the Fn promoter enriched by STAT3 binding was evaluated by agarose gel electrophoresis **(B)** or real-time quantitative PCR **(C)**; n=6 in each group. **D and E,** Immunoblots for STAT3 and p-STAT3 in the border zone (BZ) and remote zone (RZ) of the heart tissue **(D).** Quantitative analyses are plotted in **E.** n=6 in each group. **F and G,** Immunoblots for STAT3 and p-STAT3 in the cytoplasm and nucleus of cardiomyocytes from sham and MI-injured (AAV-mediated CTNT-Vector, CTNT-shSENP1, and CTNT-SENP1) mouse heart **(F).** Quantitative analyses are plotted in **G.** n=6 in each group. **H and I,** Immunofluorescence staining for p-STAT3 (red) and CTNT (green) cardiac sections; scale bar, 20 μm **(H),** Quantification of p-STAT3^+^ areas in CTNT^+^ cardiomyocyte nuclei per field (I). n=6 in each group. **J,** Immunofluorescence staining for α-SMA^+^ (red) primary fibroblasts, and S100a4^+^ (green) and α-SMA^+^ (red) NIH-3T3 fibroblasts; scale bar, 20 μm (top) and 50 μm (bottom). **K,** Quantification of α-SMA^+^ cells per field and S100a4-α-SMA double positive cells per field; n=10 in each group. One-way ANOVA with Tukey’s multiple comparisons test was conducted in **C, G, I,** and **K.** One-way ANOVA followed by the Dunn post hoc multiple comparisons test was conducted in **E.**

Recent studies have shown that overexpression of SUMO2 promotes STAT3 phosphorylation,^16^ and another study found that SUMOylation of the STAT3 Lys451 site promotes binding to the phosphatase TC45.^17^ To determine whether SENP1 directly regulates the sustained activation of STAT3, we combined the STAT3 Lys451 mutation on a SENP1-deficient background and we found that mutation of the SUMOylation site of STAT3 failed to inhibit the sustained phosphorylation of STAT3 induced by SENP1 deletion (Figure S8B and S8C). Furthermore, mutation of STAT3 Lys451 did not affect the binding of STAT3 to the *Fn* promoter (Figure S8D), suggesting that SUMOylation at the STAT3 Lys451 site failed to affect STAT3-mediated transcription of Fn.

### SENP1 Interacts with HSP90ab1 and Regulates Its Stability

To investigate the molecular mechanism of SENP1 regulating STAT3 signaling in cardiac myocytes, we predicted the high-priority proteins interacting with STAT3 using the STRING database (https://cn.string-db.org/), including EGFR, JAK1, SRC, HSP90, and EP300 (Figure S9A). We then performed immunoprecipitation-mass spectrometry with anti-SENP1 and anti-SUMO1 in primary cardiomyocytes (Online Table V). We did not find any high-priority proteins binding to SENP1 or SUMO1 by mass spectrometry, except for members of the HSP90 family. Among the interacting proteins of SENP1 and SUMO1, we identified HSP90ab1 (Figure 7A), which is a key regulator of protein homeostasis under both physiological and pathological conditions in adult cardiomyocytes and binds to STAT3 as a molecular chaperone.^18, 19^ Given the accumulation of STAT3 in the nucleus upon SENP1 deletion, this interaction is particularly interesting. Coimmunoprecipitation analysis showed that endogenous HSP90ab1-SUMO1 and HSP90ab1-pSTAT3 interactions were significantly increased in the shSENP1 group compared to controls in cardiomyocytes, which was reversed by overexpression of SENP1 (Figure 7B). The interaction between HSP90ab1 and SENP1 was then confirmed by coimmunoprecipitation of overexpressed Flag-HSP90ab1 and His-SENP1 in HEK-293T (human embryonic kidney) cells. Indeed, we found that SUMO1 could directly interact with HSP90ab1 and that this interaction was reduced after SENP1 overexpression (Figure 7C and 7D). We predicted three SUMOylation sites of HSP90ab1 at K72, K186, and K559 using JASSA (http://www.jassa.fr/index.php?m=jassa) (Figure 7E). Targeting analysis of the mutation of the three lysine residues to arginine on HSP90ab1 (HSP90ab1 K72R, HSP90ab1 K186R or HSP90ab1 K559R) showed that HSP90ab1 K72 was the most critical SUMOylation site (Figure 7F). To determine the effect of SENP1 on HSP90ab1 in cardiomyocytes, we restricted the expression of SENP1 in NRVMs. We found that SENP1 inhibition did not affect the mRNA level of HSP90ab1 (Figure S9B), and further analysis of single-cell sequencing data showed that damaged cardiomyocyte cluster did not alter HSP90ab1 gene expression (Figure S9C). However, SENP1 deletion inhibited cycloheximide-induced HSP90ab1 degradation (Figure 7G). We therefore hypothesized that SENP1 might promote HSP90ab1 ubiquitination by blocking HSP90ab1 SUMOylation. To test this hypothesis, we further examined the degree of ubiquitination of HSP90ab1. Overexpression of SUMO1 reduced the ubiquitination level of HSP90ab1, which was reversed by mutations at the K72 site of HSP90ab1 (Figure 7H).

**Figure 7.**
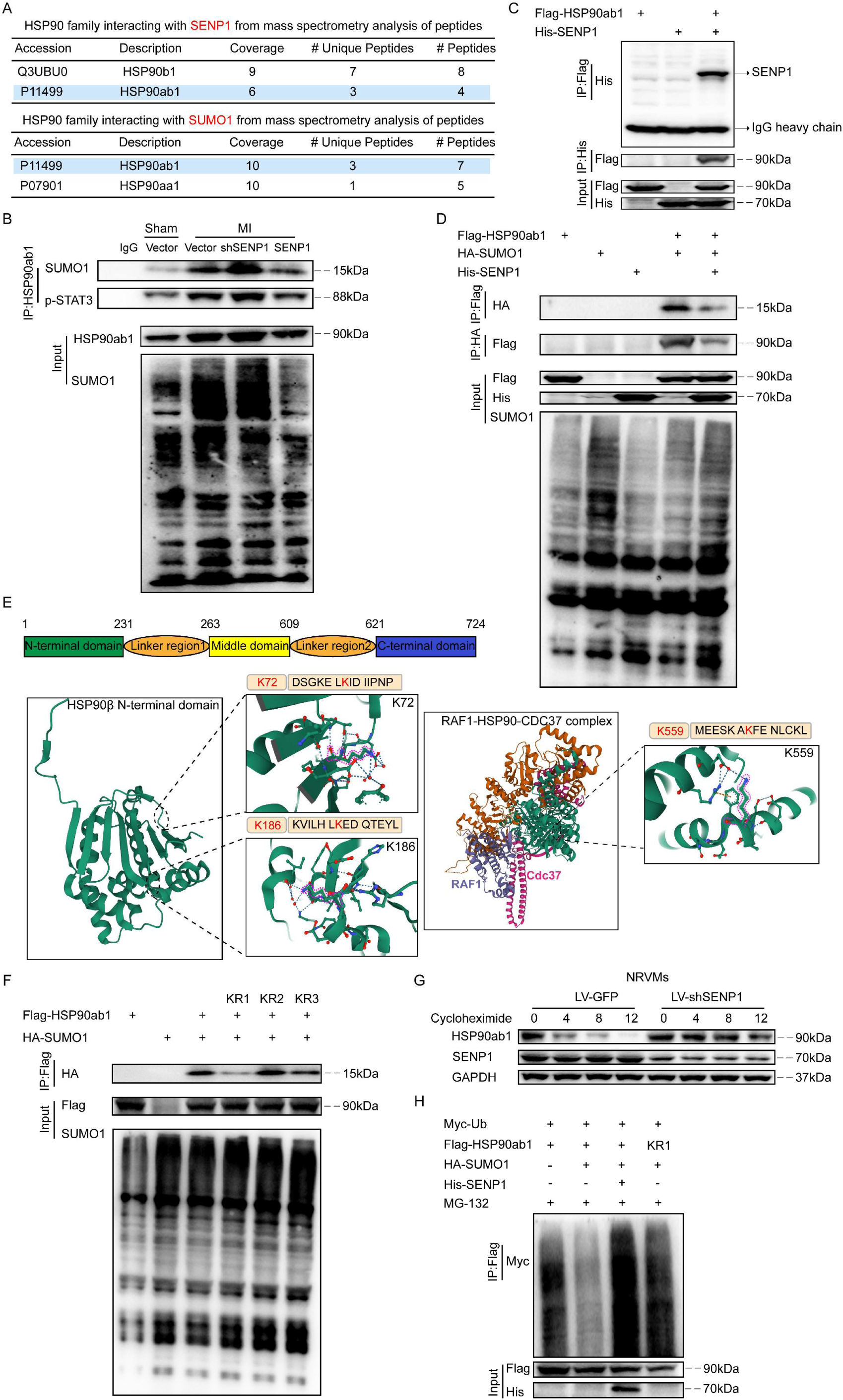
SENP1 promotes HSP90ab1 proteasome degradation by inhibiting SUMOylation at the K72 site of HSP90ab1. **A,** HSP90 family proteins interacting with SENP1 or SUMO1 from mass spectrometry analysis of peptides pulled down using anti-SENP1 or anti-SUMO1 antibody in primary cardiomyocytes. **B,** Coimmunoprecipitation (Co-IP) assays of the interaction between HSP90ab1 and SUMO1 or p-STAT3 in primary cardiomyocytes from mice treated with the indicated conditions. **C and D,** Co-IP assays of the interaction between HSP90ab1 and SENP1 **(C),** HSP90ab1 and SENP1 or SUMO1 **(D)** in HEK293T cells transfected with the indicated plasmids. **E,** The domain structure of HSP90ab1 and the K72, K186, and K559 sites in HSP90ab1 are shown. **F,** Co-IP assays using wild-type HSP90ab1 and K72(KR1), K186(KR2), or K559(KR3) mutant HSP90ab1. **G,** Immunoblotting analysis of HSP90ab1 in neonatal rat ventricular myocytes (NRVMs) infected with LV-GFP and LV-shSENP1 for 24 h and cycloheximide (CHX, 50 μM) for the indicated time points**. H,** Results of ubiquitination assays confirming the ubiquitination of HSP90ab1 after transfected with the indicated plasmids for 24 h and treated with MG132 (50 μM) for 6 h in HEK293T cells.

### Blocking HSP90 Lys^72^ SUMOylation Restored Cardioprotective Signaling and Reversing Pathological Remodeling

Because SENP1 was reduced after MI and activation of SENP1 can inhibit excessive STAT3 activation and levels in cardiomyocyte induced by HSP90ab1 over-SUMOylation, we then examined whether the SUMOylation site of HSP90ab1 could be used to treat MI-induced cardiac dysfunction. Recombinant adeno-associated virus (serotype 9) vectors carrying a mouse sumo site-mut of HSP90ab1 (HSP90ab1K72R) (accession number NM_008302.3) with a c-TNT promoter (AAV9-CTNT-HSP90ab1KR) were used to investigate the effect of SUMOylation at HSP90ab1 K72 on cardiac function after MI (Figure 8A, Figure S10A). Mice carrying HSP90ab1K72R showed preserved cardiac function versus deteriorating cardiac function in the vector group (Figure 8B and 8C). Mutation at the SUMOylation site of HSP90 significantly reduced fibrosis (Figure 8D and 8E) and suppressed Fn expression in cardiomyocytes surrounding the infarct induced by ischemic injury (Figure 8F and 8G). Furthermore, SUMOylation site depletion of HSP90ab1 abolished the effect of SENP1 deficiency on the phosphorylation and accumulation of STAT3 in the nucleus of cardiomyocytes (Figure 8H and 8I). In addition, mutations at the K72 site of HSP90ab1 in cardiomyocytes inhibited the strong pro-fibrotic effect mediated by SENP1 deletion (Figure S10B and S10C). Taken together, these data demonstrate that HSP90ab1 serves as an essential target of SENP1 in myocardial ischemia and that targeting HSP90ab1 SUMOylation may provide therapeutic strategies for pathological cardiac remodeling (Figure 8J).

**Figure 8.**
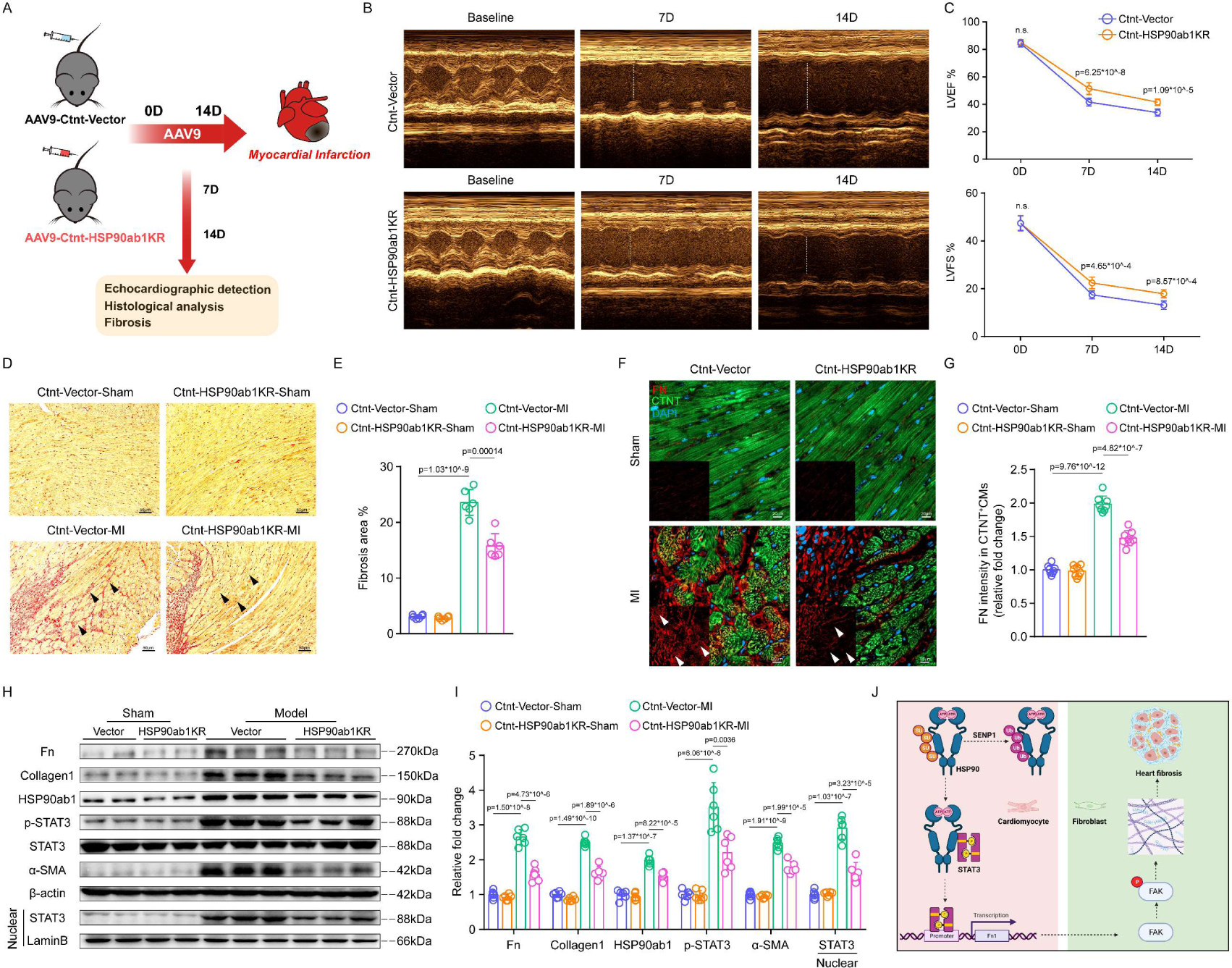
Cardiomyocyte-specific HSP90ab1 K72R gene therapy attenuates MI– induced cardiac dysfunction and fibrosis. **A,** Schematic of MI model and echocardiography in CTNT-Vector mice and CTNT-HSP90ab1KR mice. **B,** Representative echocardiographic images of CTNT-Vector mice and CTNT-HSP90ab1KR mice after MI. **C,** LVEF and LVFS measured by 2-dimensional echocardiography on mice, at different time points (0, 7, and 14 days after MI); n=8 in each group. **D and E,** Representative images of left ventricle stained with Sirius red staining showing degrees of fibrosis in mice; scale bar, 50 μm **(D).** Quantitative analyses are plotted in **E.** n=6 in each group. **F and G,** Immunofluorescence staining for Fn^+^ (red) and CTNT^+^ (green) cardiac sections; scale bar, 20 μm **(F).** Quantitative analyses of Fn^+^ areas in CTNT^+^ cardiomyocytes per field are plotted in **G;** n=8 in each group. **H and I,** Immunoblots for Fn, collagen 1, HSP90ab1, STAT3, p-STAT3 and α-SMA, in primary cardiomyocytes from mice **(H).** Quantitative analyses are plotted in **I.** n=6 in each group. **J,** Graphic illustration. In cardiomyocytes, HSP90ab1 is SUMOylated at lysine 72 and resists ubiquitination after MI, followed by STAT3-mediated activation of Fn transcription and secretion, thereby promoting FKA signaling-dependent fibroblast proliferation and activation. Cardiomyocyte-specific SENP1 overexpression or HSP90ab1K72 mutation reduces its stability and exerts cardioprotective effect after MI. Two-way ANOVA followed by Sidak post hoc multiple comparisons test was conducted in **C, E, J,** and **I.**

## Discussion

Pathological ventricular remodeling is a major component of ischemic heart disease. An incomplete understanding of the molecular mechanisms of cardiac fibrosis after MI has hindered the development of effective strategies to reverse heart failure. In this study, we report, for the first time, that SENP1 is decreased develops 5-7 days after MI, a critical time window when the adaptive/pathological remodeling transition occurs, and it is a key endogenous regulator of myocardial homeostasis in the physiological and pathological heart. We found increased levels of SENP1 14 days after MI (Figure 1). We speculate that this may be due to the massive proliferation and recruitment of fibroblasts, which is also reflected in the single cell data. Specifically, conditionally deleted SENP1 in adult cardiomyocytes induces impaired cardiac function, and exacerbates the adverse cardiac remodeling response after MI. Conversely, increasing SENP1 abundance in cardiomyocytes inhibits Fn secretion and fibroblast activation, suggesting that SENP1 is required for maintaining cardiomyocyte homeostasis and is protective against stresses altering cardiomyocyte proteostasis.

In this study, we made 3 novel observations, First, we observe that SENP1 deficiency in cardiomyocytes promotes Fn secretion and induces progressive myocardial fibrotic responses. Expression of the ECM protein Fn maintains at high levels during early postnatal development and declines in adulthood. After a pathological injury such as myocardial ischemia or pressure overload, Fn is re-induced in the myocardium and exerts paracrine effects to mediate the ventricular remodeling response. Additionally, epicardial-secreted Fn mediates epicardial-cardiomyocyte crosstalk and drives the maturation of hPSC-derived cardiomyocytes.^20^ Previous studies have shown that Fn stimulates collagen production in renal epithelial cells in a dose-dependent manner,^21^ and similarly, our study shows that Fn administration promotes fibroblast activation by mediating the activation of integrin-FAK signaling. Therefore, sustained Fn production and deposition in the extracellular matrix increases cardiac stiffness while continuously stimulating fibroblast-to-myofibroblast conversion and increased collagen secretion, leading to accelerated deterioration of adverse remodeling after MI. Genetic deletion^22^ or pharmacological suppression^14^ of Fn delays myocardial hypertrophy in pressure overload or ischemic cardiac remodeling, thereby improving short-term survival. Although Fn has been extensively studied in the autocrine regulation of fibroblasts,^23–, 25^ the regulatory mechanisms and paracrine roles of Fn in cardiac myocytes have been rarely been reported. In our study, we found that cardiomyocytes around the infarct area significantly expressed and released Fn to mediate cardiomyocyte-fibroblast communication, which was also reflected in our single-cell sequencing dataset. We determine that Fn was highly expressed in cardiomyocyte subsets after MI, especially in the Nppa^+^ cluster, and thus, the stress cardiomyocytes induced by ischemic injury are responsible for the release of Fn. In fact, injured cardiomyocytes regulate surrounding cells through a variety of factors, for example, hypoxia-induced mitogenic factor (HIMF)-mediated IL-6 secretion from cardiomyocytes promotes fibroblast activation via MAPK and CAMKII signaling.^26^ Thymosin β4 (TMSB4) and prothymosin α (PTMA) are transcriptionally regulated by ZEB2 in stressed cardiomyocytes and subsequently secreted to mediate cross-talk between cardiomyocytes and endothelial cells.^27^ Although our study shows that Fn secreted by cardiomyocytes regulates myocardial fibrosis 5-7 days after MI, it is worth noting that myocardial injury is a phased process, and the paracrine effect of cardiomyocytes on inflammatory cells (including neutrophil infiltration, macrophage transformation) at the initial stage also needs to be determined in the future.

Further analysis reveals that activation and high nuclear expression of STA3 in cardiomyocytes after myocardial ischemia is responsible for high transcript levels of Fn, which is more obvious after cardiomyocyte-specific deletion of SENP1. STAT3 is a signaling molecule capable of transporting extracellular and cytoplasmic stimuli to the nucleus in response to a variety of pathological stress responses. Recent studies have extensively demonstrated the role of STAT3 in cardiac hypertrophy,^28^ ischemia/reperfusion injury,^29^ diabetic cardiomyopathy,^30^ and heart failure.^31^ Sustained Tyr705 site phosphorylation-mediated STAT3 activation and nuclear translocation is associated with poor prognosis in cardiac hypertrophy and heart failure. It has been shown that SUMOylation at the Lys451 site promotes STAT3 binding to the phosphatase TC45 and affects nuclear translocation in head and neck cancer cell lines.^17^ In contrast, we found that mutation at the Lys451 site failed to inhibit the strong phosphorylation of STAT3-Tyr705 induced by SENP1 deletion, suggesting that the reduction of SENP1 induced by ischemic injury mediates STAT3 activation and nuclear translocation is not dependent on SUMO molecule-mediated binding of STAT3 to the phosphatase TC45, and that regulation of STAT3 by SENP1 is not solely dependent on de-SUMOylation.

Second, we demonstrate that SENP1-dependent de-SUMOylation of HSP90ab1 followed by ubiquitin-mediated degradation inhibits continued activation of STAT3. We identify HSP90ab1 of the HSP90 family as a key molecule in the SNEP1-mediated regulation of STAT3 through protein database and mass spectrometry analysis. Hsp90 interacts primarily with proteins involved in transcriptional regulation and signaling transduction pathways, such as protein kinases, ribonucleoproteins, and TFS. A study shows that XL888 (an HSP90 inhibitor without affecting other kinases) inhibits the formation of HSP90-STAT3 complexes, thereby reducing the expression levels of STAT3 and p-STAT3 in early-stage hepatocellular carcinoma.^32^ In addition, luteolin blocks the association of HSP90 with STAT and mediates the degradation of phosphorylated STAT3 (Tyr705 and Ser727) via a proteasome dependent pathway.^33^ A recent study shows that the N-terminal ATP-binding pocket of HSP90 is dispensable, the compound NCT-80 degrades STAT3 by binding to the ATP-binding pocket of HSP90 and thereby disrupting the interaction of HSP90 with STAT3.^34^ The highly conserved N-terminal domain of HSP90ab1 primarily provides sites for ATP binding. In particular, numerous studies have shown that the N-terminal domain of HSP90 provides binding sites for co-chaperones, including CDC37,^35^ AHA1,^36^ and p23/Sba1,^37^ thereby regulating HSP90 activity or the ability to bind to other ligand proteins. Therefore, the amino acid sites in the N-terminal domain of HSP90 are critical for the function of HSP90. Indeed, the binding of HSP90 to SUMO1 has been demonstrated in plasma cells^38^ and neurons.^39^ However, the regulatory function of SUMO modifications on HSP90 is unclear. We predicted three SUMOylation sites (Lys72, Lys186, and Lys559.) for HSP90ab1. Of these, the Lys72 site in the N-terminal domain and Lys559 in the middle domain responded to the SUMOylation of HSP90ab1, and mutation of the Lys 72 site to Arg maximally eliminated the degree of SUMOylation of HSP90ab1.

We found that SUMOylation at the Lys 72 site resists ubiquitination modification and increases the stability of HSP90ab1. Our identification of this critical lysine site on HSP90ab1 not only advances our understanding of SENP1-HSP90ab1-STAT3 axis-mediated signaling, but may also aid in the development of future precision therapies. Finally, and most importantly, virus-mediated in vivo injection of a cardiomyocyte-specific mutation at the HSP90ab1K72 site inhibits the extent of myocardial fibrosis and improves cardiac function after MI. In fact, in addition to SUMOylation, HSP90 is regulated by a variety of post-translational modifications and is involved in cardiac disease processes. For example, S-nitrosylation of HSP90 at Cys589 promotes interaction with glycogen synthase kinase 3β (GSK3β) and increases GSK3β phosphorylation, which contributes to the process of myocardial hypertrophy.^40^ Genetic or pharmacological inhibition of HSP90 S-nitrosylation attenuates myocardial fibrosis and adverse ventricular remodeling responses.^41^ Hyper-acetylation of HSP90 inhibits the formation of the HSP90-CX43-Tom20 complex, thereby mediating mitochondrial damage in cardiomyocytes.^42^ In addition, Hsp90 is methylated by the lysine methyltransferase Smyd2 and stabilizes the scaffolding protein titin to regulate sarcomere assembly.^43^ Recent studies have shown that inhibition of HSP90 mediated by small molecule inhibitors or gene deletion improves cardiac function in myocardial ischemia,^44^ chronic heart failure,^45^ and heart aging.^46^ However, pharmacological inhibition of HSP90, including 17-AAG, geldanamycin, 17-DMAG, and celastrol, is extensive, affects multiple signaling pathways and results in low clinical conversion. Therefore, drug design based on HSP90ab1 Lys72 will be possible for the most effective treatment of post-ischemic HF.

In summary, we provide evidence that, SENP1, by promoting HSP90 degradation, prevents ischemia–mediated myocardial fibrosis. Blocking HSP90ab1 K72-SUMOylation by a cardiomyocyte-targeted gene intervention strategy may be an effective novel approach to reverse post-MI remodeling and attenuate HF progression.

## Non-standard Abbreviations and Acronyms

α-SMA: α-smooth muscle actin
CTNT: cardiac troponin T
ECM: extracellular matrix
EF: ejection fraction
FAKs: focal adhesion kinases
FS: fraction shortening
HSP90: heat shock protein 90
MI: myocardial infarction
MYH6: myosin heavy chain 6
POSTN: periostin
SENP: sentrin-specific protease
STAT3: signal transducer and activator of transcription 3
SUMO: small ubiquitin-like modifier

## Acknowledgments

The authors thank Zhiqiang Liu, PhD (TianJin Medical University) for provided SENP1^flox/flox^ mice.

## Supplemental Material

Supplemental Methods

Figure S1–S10

Tables I–V

References #47–#49

## Novelty and Significance

### What Is Known?

- After myocardial infarction (MI), the heart undergoes a pathological remodeling process accompanied by an adverse fibrotic response.
- SENP1 mediates intracellular SUMOylation homeostasis.
- Most studies examining SENP1 in cardiovascular disease only focused on the whole heart, and the role of SENP1 in cardiomyocytes after MI remains unknown.

### What New Information Does This Article Contribute?

- Cardiac expression of SENP1 was downregulated after MI injury, primarily in cardiomyocytes.
- Cardiomyocyte-specific SENP1 deletion promotes fibrotic responses in both physiologically and pathologically.
- SENP1 deletion promotes Fn expression through transcriptional regulation mediated by HSP90ab1-STAT3 in cardiomyocytes.

Adverse ventricular remodeling is a major challenge for patients after myocardial infarction (MI) and its therapeutic intervention has been unsuccessful. SENP1 is involved in the pathogenesis of cardiovascular diseases, including myocardial hypertrophy and heart failure. However, the role of cell type-specific SENP1 in the setting of MI is unclear. Here, we found that SENP1 deficiency in cardiomyocytes induces cardiac fibrosis by promoting Fn transcription and secretion. On the other hand, we found that the HSP90 Lys72 SUMOylation is key for promoting STAT3-mediated Fn transcription. Our findings suggest that blocking HSP90ab1 SUMOylation at the Lys72 site may be an effective and novel target for reversing adverse ventricular remodeling after MI.

